# Surface-Based vs. Voxel-Based Finite Element Head Models: Comparative Analyses of Strain Responses

**DOI:** 10.1101/2024.09.04.611159

**Authors:** Zhou Zhou, Xiaogai Li, Svein Kleiven

**Affiliations:** Division of Neuronic Engineering, KTH Royal Institute of Technology, Stockholm, 14152, Sweden

**Keywords:** Traumatic brain injury, finite element head models, hexahedral mesh techniques, surface- and voxel-based meshing, brain strain

## Abstract

Finite element (FE) models of the human head are important injury assessment tools but developing a high-quality, hexahedral-meshed FE head model without compromising geometric accuracy is a challenging task. Important brain features, such as the cortical folds and ventricles, were captured only in a handful of FE head models that were primarily developed from two meshing techniques, i.e., surface-based meshing with conforming elements to capture the interfacial boundaries and voxel-based meshing by converting the segmented voxels into elements with and without mesh smoothing. Despite these advancements, little knowledge existed of how similar the strain responses were between surface- and voxel-based FE head models. This study uniquely addressed this gap by presenting three anatomically detailed models - a surface-based model with conforming meshes to capture the cortical folds-subarachnoid cerebrospinal fluid and brain-ventricle interfaces, and two voxel-based models (with and without mesh smoothing) - derived from the same imaging dataset. All numerical settings in the three models were exactly the same, except for the meshes. These three models were employed to simulate head impacts. The results showed that, when calculating commonly used injury metrics, including the percentile strains below the maximum (e.g., 99 percentile strain) and the volume of brain element with the strain over certain thresholds, the responses of the three models were virtually identical. Different strain patterns existed between the surface- and the voxel-based models at the interfacial boundary (e.g., sulci and gyri in the cortex, regions adjacent to the falx and tentorium) with strain differences exceeding 0.1, but remarkable similarities were noted at the non-interfacial region. The mesh smoothing procedure marginally reduced the strain discrepancies between the voxel- and surface-based model. This study yielded new quantitative insights into the general similarity in the strain responses between the surface- and voxel-based FE head models and underscored that caution should be exercised when using the strain at the interface to predict injury.

## Introduction

Traumatic brain injury (TBI) represents a critical public health problem. Globally, around 69 million people suffer a TBI each year (Dewan et al. 2018). In Europe, TBI is the leading cause of death in young adults and its incidence is on the rise in the elderly (Brazinova et al. 2021). TBI is considered a ‘silent epidemic’, as the society largely unrecognizes the magnitude of this problem (Maas et al. 2022). In the European Union, about 7.7 million people are living with a TBI-induced disability (Peeters et al. 2015). The economic cost of TBI is enormous with a yearly cost of around 33 billion Euros at the European level (Olesen et al. 2012) and 400 billion Euros at the global level (Maas et al. 2017). To address this TBI-related urgency, advancing fundamental understandings of the brain injury mechanics is crucial for the development of more effective prevention strategies.

Finite element (FE) models of the human head are instrumental in providing quantitative insight into how the external impact instigates localized tissue straining and triggers the injury occurrence (Ji et al. 2022; Kleiven 2002). Owing to the intricate complexity of the human head, developing an FE head model for TBI investigation is not a trivial task with manifold variables affecting the injury prediction outcome. As reviewed by Yang and Mao (2019), the model should have sufficient representations of head sub-components (e.g., brain, ventricle) in terms of geometry (Cloots et al. 2008; Giudice et al. 2023; Huynh et al. 2024; Kleiven and Von Holst 2002; Li 2021) and material (Giordano et al. 2014; Ho et al. 2017; Li et al. 2017; Zhao et al. 2018) as well as accurate descriptions of intracranial interfaces (e.g., brain-ventricle and brain-skull interactions) (Rycman et al. 2024; Wang et al. 2018; Zhou et al. 2019a; Zhou et al. 2020b). Careful considerations should also be given to the strategy of geometrical discretization (e.g., mesh type, size, and quality) (Mao et al. 2013a; Panzer et al. 2013; Zhao and Ji 2019a), element formulation (Giudice et al. 2019; Zhao and Ji 2019a), and other numerical choices, e.g., hourglass control (Giudice et al. 2019; Takhounts et al. 2003). Before its usage of injury assessment, the model needs to be extensively evaluated by injury-relevant experimental data (e.g., brain motion, strain, and strain rate with the impact severity close to the traumatic level) (Alshareef et al. 2018; Alshareef et al. 2020; Dutrisac et al. 2021; Hardy et al. 2001; Hardy et al. 2007; Zhou et al. 2019b; Zhou et al. 2018; Zhou et al. 2023). Even for injury assessment itself, one critical challenge is how best to interpret the spatiotemporally detailed, multiform responses yielded by the FE model [e.g., maximum principal strain (MPS) in the whole brain vs. maximum von Mises stress in the corpus callosum (CC)] to achieve the optimal predictability (Fahlstedt et al. 2022; Zhao and Ji 2019b; Zhou et al. 2021; Zhou et al. 2024). Thanks to the endeavour of the TBI research community over the past decades, modern FE head models are featured with subject-specific geometry (Giudice et al. 2020; Li 2021), hyper-viscoelastic brain material with tension-compression asymmetry and rate dependency (Ji et al. 2015; Kleiven 2007), and solid and fluid representations of the intracranial components (Zhou et al. 2019a; Zhou et al. 2020b). Encouraging correlations have been reported between the FE-derived mechanical loading at the brain tissue or its microstructures and clinically diagnosed injury in real life (e.g., concussion, parenchymal haemorrhage) (Fahlstedt et al. 2015; Montanino et al. 2021; Yuan et al. 2024; Zhang et al. 2024).

Two important features of the human head are the cortical folds (i.e., gyri and sulci) and ventricles, both of which have important biomechanical roles in the brain’s vulnerability. For example, by virtually impacting two FE head models with and without sulci, Ho and Kleiven (2009) found that high strain level was concentrated to several spots at the depth of the sulci, as was laterally confirmed in another FE study (Ghajari et al. 2017). Such a strain distribution largely overlapped with the site of certain brain pathologies, e.g., cerebral infarcts in diffuse axonal injury (Strich 1956) and tau pathology in chronic traumatic encephalopathy (McKee et al. 2013). Recently, Zhou et al. (2020a) reported the presence of the ventricles, a cerebrospinal fluid (CSF)-filled cavity, exacerbated the strain in the deep brain regions (e.g., ventricular wall, CC, thalamus), providing a possible explanation for the prevalence of periventricular injury in severe TBI patients. Despite their biomechanical prominence, the cortical folds and ventricles were largely simplified in the existing FE head models (Garimella and Kraft 2017; Ji et al. 2015; Kimpara et al. 2006; Kleiven 2007; Mao et al. 2013b; Takhounts et al. 2008b; Trotta et al. 2020; Zhang et al. 2001; Zhou et al. 2016), of which the brain was simplified by smearing out the gyri and sulci with a homogeneous layer of subarachnoid CSF, and certain elements were selected to represent the ventricles.

Generating high-quality meshes without compromising geometric and morphologic details is a challenging task. This is especially the case for the brain as the hexahedral element is generally preferred owing to its stability and accuracy in scenarios with large deformation (e.g., TBI modelling) and material incompressibility (e.g., brain and CSF) (Giudice et al. 2019). Among the handful of pure hexahedral FE head models featuring cortical folds and ventricles, two different meshing techniques were used. The first one was a surface-based meshing, in which outer surfaces of individual components (e.g., pial surface, ventricular surface) were first generated and hexahedral meshes were deployed to fill the domain encased within the inputted surfaces and capture the boundaries between different components (Li et al. 2017; Li et al. 2019; Li et al. 2021; Zhou et al. 2022a; Zhou et al. 2020a). The other prevalently used method was a voxel-based meshing, in which segmented voxels were converted into hexahedral elements (Alshareef et al. 2021; Chen and Ostoja-Starzewski 2010; Duckworth et al. 2022; Giudice et al. 2020; Griffiths et al. 2023; Ho and Kleiven 2009; Ho et al. 2009; Ho et al. 2017; Miller et al. 2016; Pavan et al. 2022). Despite the success of both techniques in generating anatomically detailed FE head models, no knowledge existed of how similar the responses were between the surface- and voxel-based models of the human brain. To the best of the authors’ knowledge, only Bradfield (2022) compared the strain responses from two rat (lissencephalic) brain models (one with surface-based meshes and the other with voxel-based meshes with perfect cubes). It was reported that the strain differences between these two models were small, but some strain discontinuities were noted at the brain-skull interface in the voxel-based model. When limited to the voxel-based model, a mesh smoothing procedure was often implemented to reduce the jaggedness between different parts (e.g., the interface between the brain and CSF) (Chen and Ostoja-Starzewski 2010; Duckworth et al. 2022; Griffiths et al. 2023; Ho and Kleiven 2009; Ho et al. 2009; Ho et al. 2017; Huynh et al. 2024; Wang et al. 2020). Although the smoothing procedure was known to reduce the mesh quality (Kelley et al. 2015), its influence on brain strain has not been quantified before.

The current study aimed to answer the research question of how close the strain responses were between the surface- and voxel-based FE head models. To this end, a previously developed surface-based head model (Li et al. 2021) was reused in this study with conforming elements to capture the cortical folds-subarachnoid CSF interface and the brain-ventricle interface, and two voxel-based models were newly created by converting voxels as elements with and without mesh smoothing, respectively. We kept all numerical settings in the three models identical, except for the meshing. This enabled a focused investigation on the effect of meshing the same head geometry via three different techniques on the brain strains. These three models were used to simulate head impacts, and the predicted strains were compared. The objectives of this study were dual: 1) assessing whether the strain responses of surface and voxel-based models were comparable, especially at the interfacial region; 2) evaluating if the mesh smoothing procedure mitigated the strain differences between the surface- and voxel-based models.

## Methods

### Mesh generation

For the surface-based meshing, its implementation was elaborated by Li et al. (2021) early with the delivered FE head model (Fig 1I) reused in the current study. In brief, the International Consortium for Brain Mapping (ICBM) 152 atlas (voxel size: 1 × 1 × 1 mm; slice thickness: 1 mm), developed from MRI data of 152 young adults (aged 18.5 - 43.5 years) (Fonov et al. 2011; Mazziotta et al. 2001), was used as geometrical sources. The atlas was segmented using an open-access software (i.e., 3D Slicer) (Fedorov et al. 2012) based on an expectation-maximization algorithm together with the spatial information provided by the probability maps to delineate the brain (with details of gyri and sulci) and CSF (including subarachnoid CSF and ventricles) (Fig 1A). The skull was created by thresholding the atlas and then removing the brain and CSF masks with manual editing. The triangulated outer surfaces of the brain, subarachnoid CSF, ventricle, and skull were extracted from the segmentation (Fig 1B) and further input to the Hexotic software (Maréchal 2009) to automatically generate all hexahedral elements (Fig 1C) using an octree-based method (Maréchal 2009; Schneiders 2000). The octree-based method recursively split an initial box (i.e., the root octant that was a perfect cube and covered the whole input profiles) into smaller boxes (i.e., descended octants that were often irregular cubes) until the descendant octants could resolve the local geometry. The recursive level was a key parameter to execute the automatic meshing function in the Hexotic software and was primarily determined by the complexity of the geometry. The current study focused on the head with particular interests in the brain-skull interface and brain-ventricle interface, a level of eight (i.e., the root octant was recursively partitioned eight times) was found to be sufficient with the resultant hexahedral mesh conforming to the inner and outer surfaces of the skull, pial surface (Fig 1C), and ventricle (Fig 1F). This model was referred to as Conf-model hereafter.

**Fig 1.**
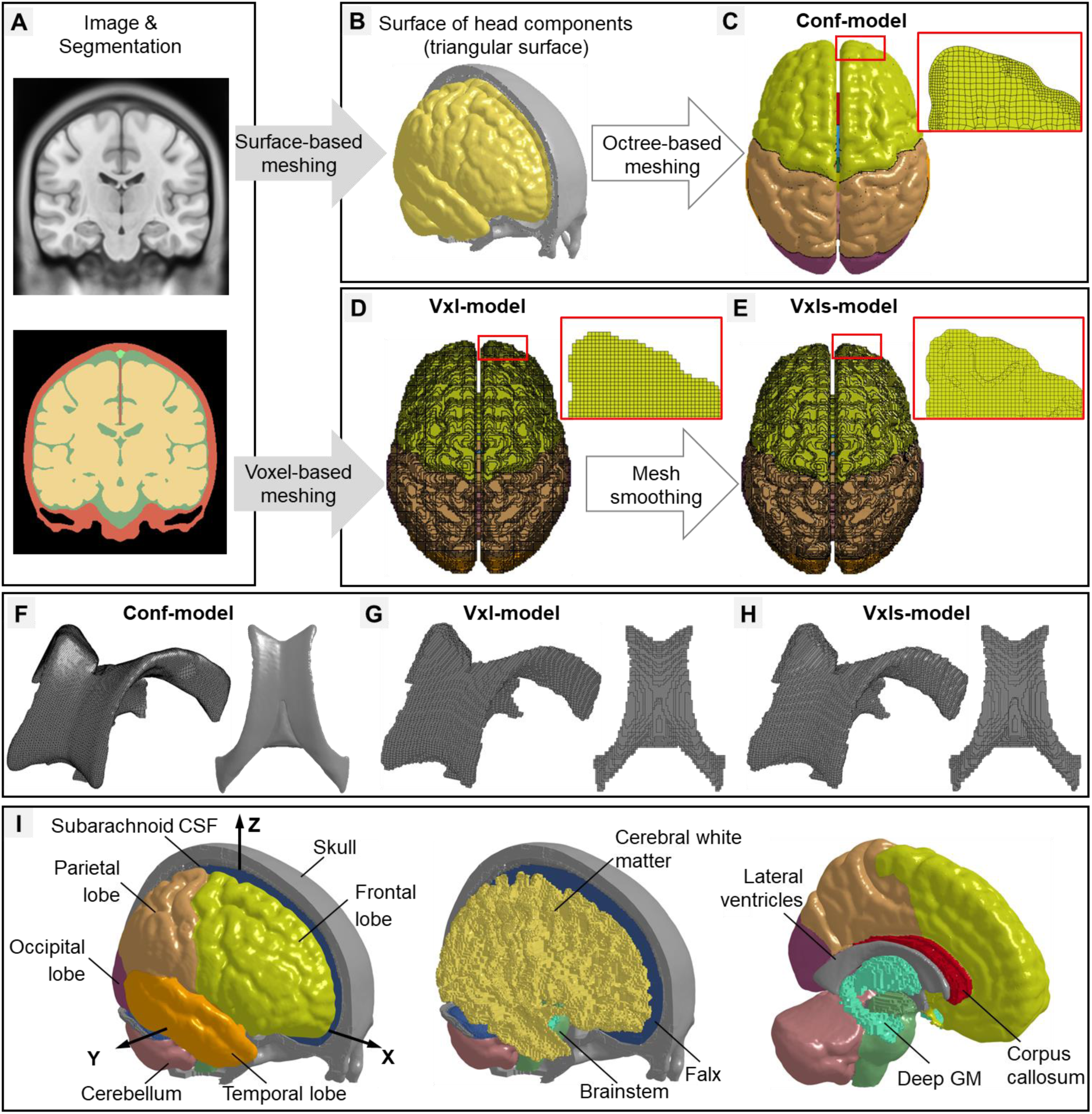
The workflow of developing three biomechanical head models with pure hexahedral meshes, named Conf-model (i.e., a model with conforming mesh for the brain-CSF interface), Vxl-model (i.e., a voxel-based model without mesh smoothing), and Vxls-model (i.e., a voxel-based model with mesh smoothing), respectively. (A) T1-weighted magnetic resonance imaging and corresponding segmentation. (B) The head geometries, i.e., skull, brain, and ventricle (unshown), were used to develop the Conf-model (C), while the segmentation was used to create the Vxl-model (D) and the Vxls-model (E). For the ventricles, an isometric view (mesh lines on) and a top view (mesh lines off) were provided for the Conf-model (F), Vxl-model (G), and Vxls-model. (H) To facilitate the post-processing, the brain elements of the three models were further grouped to have explicit representations of the major components (as exemplified by the Conf-model in subfigure I), including cerebral gray matter (GM) (i.e., frontal, parietal, occipital, and temporal lobes), cerebral white matter, deep GM (i.e., hippocampus and thalamus), corpus callosum, cerebellum, and brainstem. A skull-fixed coordinate system (XYZ, X: posterior-anterior; Y: left-right; Z: inferior-superior) was illustrated with the origin at head’s center of gravity.

For the voxel-based meshing, a head model (referred to as Vxl-model) was newly developed by directly converting each isotropic voxel in the ICBM atlas (Fig 1A) into a cubic hexahedral element (Fig 1D). Elements with their voxel counterparts that have the same segmentation label were assigned to one part (i.e., brain, subarachnoid CSF, ventricle, skull), the same as the approaches used to develop other models (Alshareef et al. 2021; Giudice et al. 2020; Miller et al. 2016; Pavan et al. 2022). Note that, in the segmentation directly output form 3D-Slicer, the brain was parcellated into many subregions, each of which was identified by a unique label, similar to the case in the study by (Huynh et al. 2024). Prior to converting the voxel to element, we combined the labels of brain subregions as one. Critically, the resultant mesh had stair-stepped interfaces between the brain and CSF, as was partially evidenced by the enlarged view of the brain surface in Fig 1D and the ventricles in Fig 1G.

To alleviate the issue of jagged interfaces, a Laplacian-based, volume-preserving mesh smoothing algorithm (Boyd and Müller 2006) was applied to the Vxl-model, thereby creating a new derivative model (referred to Vxls-model, Fig 1E). This smoothing algorithm was implemented via an in-house customized script. In brief, it optimized the locations of those nodes at the exterior surface (i.e., outer surface of the skull) and intracranial interfaces (e.g., skull-CSF interface, CSF-brain interface, and brain-ventricle interface) to balance the need of reducing the jaggedness level between different tissues (see the enlarged view of the brain surface in Fig 1E and the ventricles in Fig 1H) and preserving the volume of head components and element quality. The derivative Vxls-model retained the same element and node numbers and mesh topology as the original Vxl-model. Technical details of this smoothing algorithm were initially presented by Boyd and Müller (2006) and its application for smoothing voxel-based FE head models was noted in earlier studies (Chen and Ostoja-Starzewski 2010; Wang et al. 2020).

In the three models, the pia mater and dura mater were additionally created as shell elements by finding the outer surfaces of the brain and subarachnoid CSF, respectively. The falx and tentorium were manually generated based on the tissue classification from in-house high-resolution brain MRI data and anatomical knowledge acquired from early conducted human head tissue experiments (Walsh et al. 2021a) and literature review (Walsh et al. 2021b). The falx and tentorium among the three models matched in size. All membranes were modelled as quadrilaterals with fully integrated Belytschoko-Tsay membrane [ELFORM = 9, in LS-DYNA (Hallquist 2007b)] and share common nodes with the neighbouring CSF. Except for stemming from the same imaging dataset for the head geometry as detailed above, the three models also shared commonalities in terms of the material property, element formulation, and interface condition, as summarized in Appendix A in the current study and also detailed in the early article by Li et al. (2021). In brief, the brain was modelled as a homogenous isotropic structure, but it was further grouped into different sub-components (Fig 1I) to facilitate postprocessing. The subarachnoid CSF and ventricles were modelled as an elastic fluid. A selectively reduced element formulation (ELFORM = 2, in LS-DYNA) was used for the brain and CSF. This formulation excelled in using reduced integration for the volumetric (pressure) term, and full integration for the deviatoric (shear) terms, circumventing the hourglass issue in the reduced integration formulation and volume locking phenomenon in the full integration formulation of lower order elements (Hallquist 2006a; Hallquist 2006b; Hallquist 2007a; Hallquist 2007c; Kleiven 2007). All intracranial interfaces (i.e., brain-skull interface, brain-ventricle interface, falx/tentorium-CSF interface) shared common nodes, a prevalently used approach in the existing FE head models (Ji et al. 2022; Wang et al. 2018). Responses of the Conf-model have been extensively evaluated by experimental data of intracranial pressure, brain-skull relative displacement, brain strain (Li et al. 2021), and brain strain rate (Zhou et al. 2023). Additional validations of the Vxl- and Vxls-models were not performed, owing to the response similarity among the three models (see the results section).

Given its importance of numerical accuracy and stability, the mesh quality of the Conf-, Vxl-, and Vxls-models was examined based on 6 indices (pass/fail criteria), i.e., Jacobian (0.5), warpage (30°), skew (60°), aspect ratio (8), maximum angle (30°), and minimal angle (150°). These indices have been commonly used to assess mesh quality in the literature (Ho et al. 2009; Kelley et al. 2015; Mao et al. 2013a; Zhou et al. 2016) and the criterion of each index was the same as those adopted by Li et al. (2021).

### Impact simulations

To compare the strain responses between the surface-based model and voxel-based models with and without smoothing, impact simulations were performed using Conf-, Vxl-, and Vxls-models. In one National Football League impact dataset, the head kinematics were obtained from laboratory reconstruction of on-field video-recorded collisions using Hybrid III anthropometric test dummies (Sanchez et al. 2019). Ten cases, covering different impact directions and severities, were selected from this dataset. The acceleration peaks were summarized in Table 1 and detailed acceleration profiles of one representative case (i.e., Case 1) were exemplified in Figure 2. For each impact, the directional linear acceleration and angular velocity curves were imposed to one node located at the head’s centre of gravity and constrained to the rigidly modelled skull. For a simulation of 60 ms, it took 8 hours for the Conf-model, 4 hours for the Vxl-model, and 5 hours for the Vxls-model, all of which were solved by the massively parallel processing version of LS-DYNA 13.0 version with 128 central processing units. The explicit central difference method was used for the time integration (Hallquist 2006b). The timesteps were 0.111 microsecond for the Conf-model, 0.585 microsecond for the Vxl-model, and 0.277 microsecond for the Vxls-model, all of which was dictated by the CSF. The model responses were output at every 0.5 millisecond.

**Fig 2.**
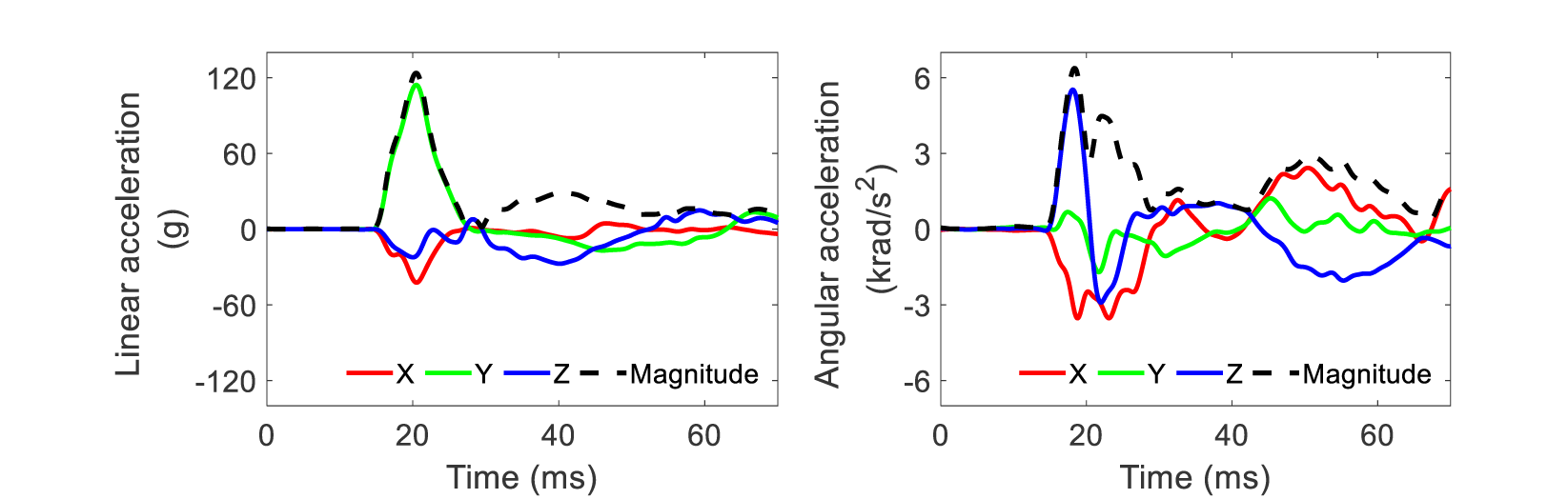
Acceleration curves for one representative impact (Case 1). Note that the X, Y, and Z axes are the same as those in the skull-fixed coordinate system in Fig 1F.

**Table 1.** Peaks of linear acceleration and angular acceleration of the 10 simulated cases. Note that the X, Y, and Z axes are the same as those in the skull-fixed coordinate system in Fig 1I.

| Case ID | Peak linear acceleration (g) |  |  |  | Peak angular acceleration (krad/s<sup>2</sup>) |  |  |  |
| --- | --- | --- | --- | --- | --- | --- | --- | --- |
|  | X | Y | Z | Magnitude | X | Y | Z | Magnitude |
| Case 1 | -42.3 | 114.3 | -27.4 | 123.7 | -3.53 | -1.70 | 5.53 | 6.37 |
| Case 2 | -29.2 | 22.5 | -34.6 | 49.4 | -2.38 | -1.01 | 1.10 | 2.77 |
| Case 3 | 19.7 | 56.3 | -6.0 | 59.8 | -4.04 | 2.95 | -6.74 | 8.30 |
| Case 4 | -21.5 | 59.4 | -57.8 | 78.8 | -5.82 | 1.66 | 2.43 | 6.23 |
| Case 5 | -31.9 | -133.3 | -41.6 | 138.2 | 4.65 | 1.35 | 6.82 | 7.50 |
| Case 6 | -78.0 | 66.5 | -21.8 | 102.1 | -3.21 | -2.39 | 4.40 | 4.98 |
| Case 7 | -66.8 | -20.0 | -42.9 | 79.2 | 2.45 | -3.44 | -1.27 | 4.29 |
| Case 8 | -64.1 | 79.9 | 44.5 | 103.2 | -5.69 | -4.70 | 3.88 | 7.54 |
| Case 9 | -26.4 | 8.4 | -13.5 | 29.3 | -0.86 | -1.41 | -0.94 | 1.53 |
| Case 10 | -31.7 | 40.0 | -18.2 | 52.3 | -2.77 | -1.87 | -2.21 | 3.13 |

### Strain analyses

We assessed the influences of meshing technique on strain responses. For each element, we calculated the MPS of as the first eigenvalue of the Green-Lagrange strain tensor at each time instance, and identified the maximum MPS value during the entire simulation (denoted as *ε*_*conf*_*^sim^* for the Conf-model, *ε_vxl_ ^sim^* for the Vxl-model, and *ε _vxls_^sim^* for the Vxls-model). Given the common suspicion that the FE head model, especially voxel-based meshing, was prone to have strain concentration (Giudice et al. 2019; Madhukar and Ostoja-Starzewski 2019; Mao et al. 2013a), the maximum MPS value across all elements (determined by a single element with the highest MPS than its counterparts) was vulnerable to be affected by numerical artifacts. Instead, percentile MPS values (e.g., 99^th^ percentile, 95^th^ percentile, 90^th^ percentile) were alternatively calculated. For example, the 95^th^ percentile MPS was mathematically computed as the 95 percentile of element-wise MPS array (assuming the elements with MPS values in the top 5 percentile had numerical instability and should be excluded). Since no consensus was reached on the choice of percentile value (Fahlstedt et al. 2022; Östh et al. 2023; Panzer et al. 2012), the strain values with the percentile ranging from 0 to 100 (‘*prctile*’ function in Matlab) were calculated. Similar percentile plots were reported in earlier studies [e.g., Garlapati et al. (2015); Wang et al. (2018)]. Note that the maximum MPS value across all elements was equivalent to the 100 percentile MPS value. In addition, the volume fraction of elements with the MPS magnitude over certain thresholds was also examined, similar to the widely used cumulative strain damage measure (Takhounts et al. 2008a). Both the magnitude- and volume-based calculations were implemented to the three models at four regions of interest (ROI), i.e., cerebral gray matter (GM) (including frontal, occipital, parietal, and temporal lobes), cerebral white matter (WM), CC, deep GM (including hippocampus and thalamus), and the whole brain level.

To further assess the similarity of strain patterns and localize the potential region with high strain differences among the three models, spatially detailed comparisons were performed based on voxelized strain results by mapping the element-wise brain MPS responses into the ICBM space (voxel resolution: 1 mm^3^, brain voxels: 1,567,589). The mapped strain peaks were denoted as *ε*_*conf*_*^map^* for the Conf-model, *ε _vxl_^map^* for the Vxl-model, and *ε _vxls_^map^* for the Vxls-model. This mapping procedure was intuitive for the Vxl-Model and Vxls-Model owing to the one-one relationship between the element and the voxel. Thereby, *ε _vxl_^map^* was identical to *ε_vxl_ ^sim^*, and *ε_vxls_^map^* was equivalent to *ε_vxls_^sim^* . For the Conf-model, the brain strain response was at the element level with spatially heterogeneous resolution, mismatching with the voxel resolution. To address this, the distances between voxels and elements were calculated. For a given voxel, its counterpart in the Conf-model was determined as the closest brain element in terms of Euclidean distance. The spatial deviations between paired voxels and elements were all less than 0.5 mm. Appendix B illustrated the strain response directly simulated by the Conf-model (*ε*_*conf*_*^sim^*) and its counterpart mapped in the imaging space (*ε*_*conf*_*^map^*). Identical distribution and magnitude was noted between *ε^sim^_conf_* and *ε^map^_conf_*, supporting the validity of the voxelization procedure. Similar approaches have been used by Ji and Zhao (2022) and Zhou et al. (2022b) to resolve mesh-voxel mismatch. Voxel-wise comparisons were performed among *ε^map^_conf_*, *ε_vxl_^map^*, and *ε^map^_vxls_* in an iterative manner. For any strain pairs, Pearson’s correlation coefficient (R) was determined, and the root-mean-square error normalized by the mean value of the independent variables (NRMSD) was reported. The threshold for significance was *p <* 0.05.

## Results

### Strain responses

For one illustrative impact (Case 1), contours of element-wise MPS results directly from the Conf-, Vxl-, and Vxls-models (i.e., *ε_conf_^sim^*, *ε_vxl_^sim^*, and *ε_vxls_ ^sim^*) were qualitatively compared in Fig 3A-I. Remarkably similar strain patterns were noted among the three models with the high strain (characterized by red colour) commonly distributed at the peripherical area. A visual scrutinization of the enlarged view showed abrupt discontinuity in strain distribution at the cortex of the Vxl-model (Fig 3F), in which elements at the brain surface had much smaller strains than their underlying neighbouring counterparts. Such a discontinuity was less prominent in the Vxls-model (Fig 3I) and further mitigated in the Conf-model (Fig 3C).

**Fig 3.**
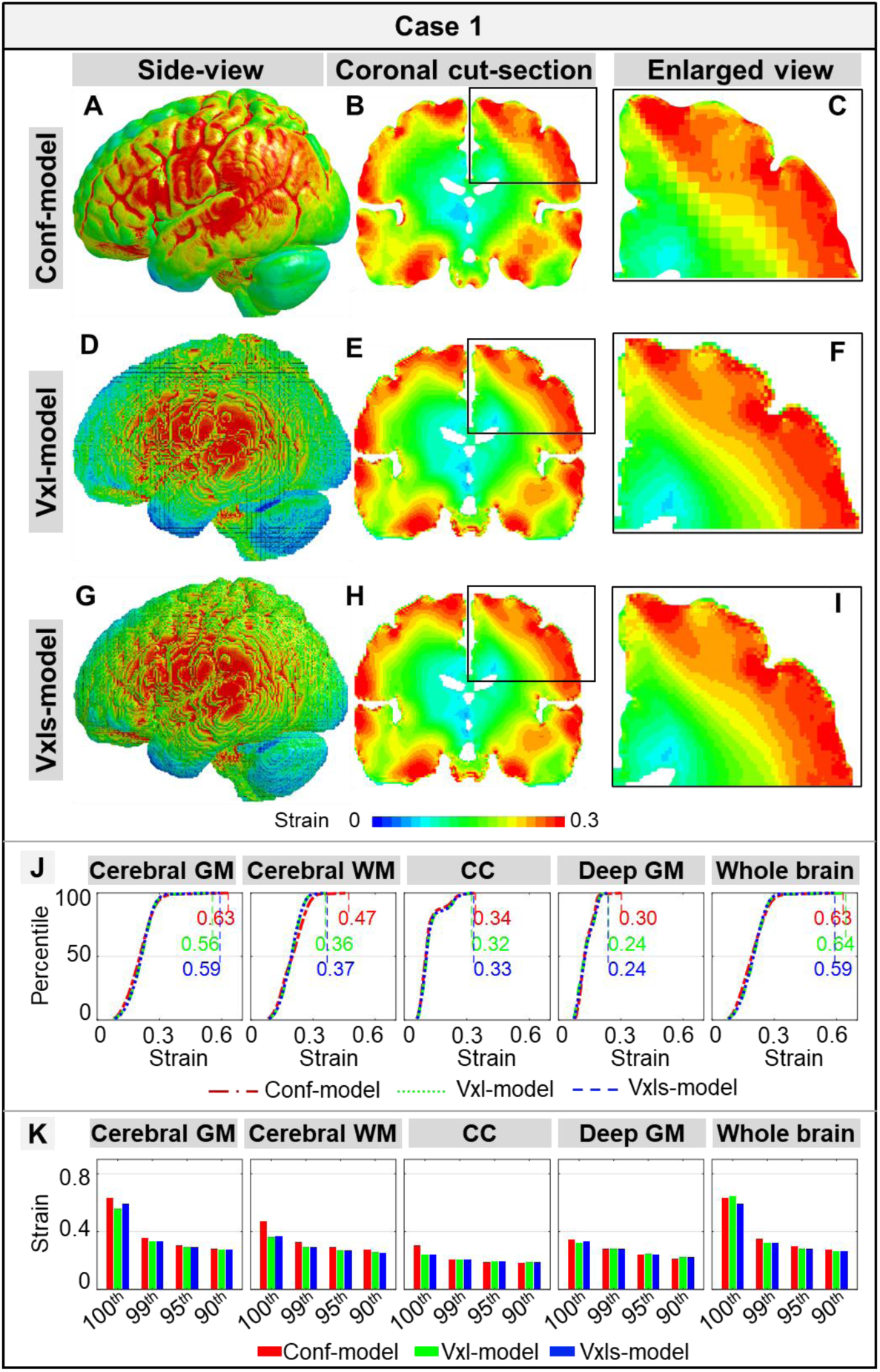
Comparisons of time-accumulated strain results among the Conf-, Vxl- and Vxls-models in Case 1. Strain results of the Conf-model were shown in the whole-brain side view (A), a coronal cut-section (B), and an enlarged view (C). Similar illustrations were presented for the Vxl-model (D-F) and Vxls-model (G-I). (J) Cumulative percentile strain curves at four regions of interest (i.e., cerebral gray matter (GM), cerebral white matter (WM), corpus callosum (CC), deep GM) and the whole brain, of which the 100^th^ percentile strain peak values were listed. (K) The strain peak values of 100^th^ percentile, 99^th^ percentile, 95^th^ percentile, and 90^th^ percentile.

For the magnitude-based strain comparisons, the cumulative percentile strain curves in Case 1 (Fig 3J) were generally overlaid with each other with exceptions noted at the 100^th^ percentile value (both for the four ROIs and the whole brain). For example, the 100^th^ percentile strain values in the cerebral GM were 0.63 for the Conf-model, 0.56 for the Vxl-model, and 0.59 for the Vxls-model. This was further confirmed in Fig 3K, in which the differences in the enumerated 99^th^, 95^th^, and 90^th^ percentile strain peaks among the three models were less than 0.02. For Cases 2-10 (Fig C1 in Appendix C), similar trends were noted among the three models, of which the cumulative percentile strain curves were superimposed on each other with the disparity commonly noted in the 100^th^ percentile value.

For volume-based strain comparisons, comparable values were noted among the three models in terms of the volume fraction of elements with the MPS value over 0.2 for Case 1 (Fig 4A). The cross-model differences were less than 4%. For example, 16.2% of CC volume in the Conf-model endured a MPS value over 0.2, in contrast to 13.0% in the Vxl-model and 12.7% in the Vxls-model. When expanding to other strain thresholds and other impacts, curves that quantified the volume fractions of elements with the MPS value over different thresholds were largely overlaid among the three models for Case 1 (Fig 4B) and Cases 2-10 (Fig C2 in Appendix C).

**Fig 4.**
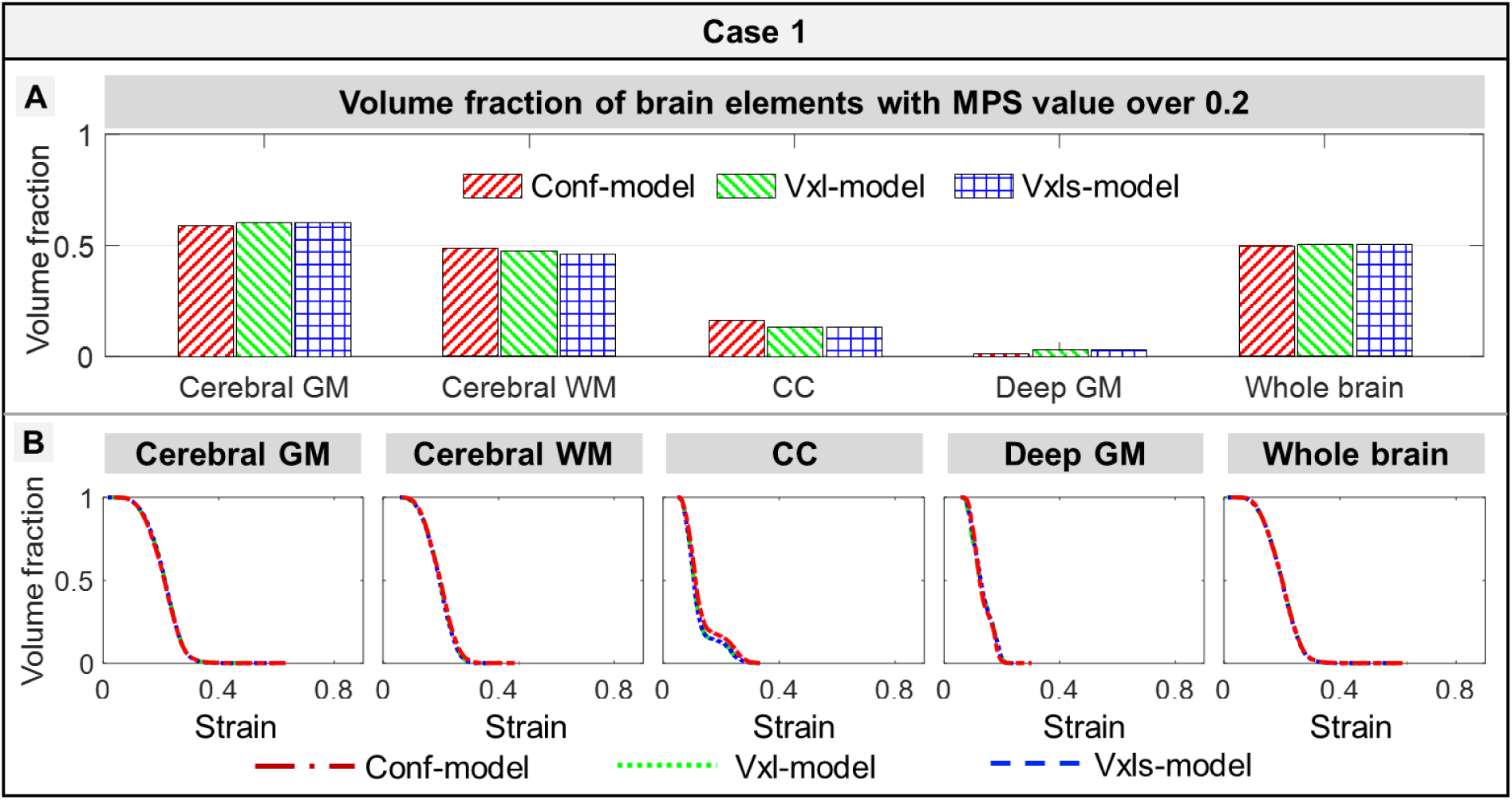
Comparisons of volume fractions of brain elements over varying threshold among the Conf-, Vxl- and Vxls-models in Case 1. (A) Volume fraction of brain elements with MPS value over 0.2 in cerebral gray matter (GM), cerebral white matter (WM), corpus callosum (CC), deep GM, and the whole brain estimated by the Conf-, Vxl- and Vxls-models. (B) Cumulative curves of volume fraction of the elements with the strain over varying thresholds. These curves were calculated in the cerebral GM, cerebral WM, CC, deep GM, and whole brain. It should be clarified that, when the threshold was chosen as the 100^th^ percentile strain of one model, the volume fractions among the three model would be different. As the resultant volume fractions were all close to or equal to zero, this threshold choice made little sense and hence was not discussed later.

When switching to the voxelized strain response (i.e., *ε*_*conf*_*^map^* for the Conf-model, *ε_vxl_^map^* for the Vxl-model, and *ε _vxls_^map^* for the Vxls-model), an illustrative comparison for Case 1 was shown in Fig 5A-F. The largest disparity was noted between *ε*_*conf*_*^map^* and *ε_vxl_ ^map^* with the R-value as 0.91 and the NRMSD as 11.7% (Fig 5A). 13,457 voxels (vs. all brain voxels: 1,567,589) had high strain differences (i.e., the absolute value of strain difference over 0.1) and were primarily located at interfacial boundaries (i.e., sulci and gyri in the cortex, regions adjacent to the falx and tentorium) in a cluster manner (Fig 5B). When switching to the strain pair of *ε*_*conf*_*^map^* and *ε_vxls_^map^*, the R-value was increased to 0.94 (NRMSD: 9.5%) (Fig 5C). The number of voxels with high strain differences was 5,092 and those voxels were primarily aggregated around the tentorium and spread the whole pial surface in a dotted manner (Fig 5D). For the correlation between *ε_vxls_^map^* and *ε_vxl_ ^map^*, the R-value was 0.98 (NRMSD: 5.4%) (Fig 5E). Only 763 voxels (primarily around the lateral sulcus) had strain differences over 0.1 (Fig 5F). When collectively scrutinizing *ε_conf_^map^*, *ε_vxl_^map^*, and *ε_vxls_^map^*, no consistent findings were noted regarding their relative magnitudes as both positive and negative values were noted in the subtraction of any strain pairs (Fig 5A-F).

**Fig 5.**
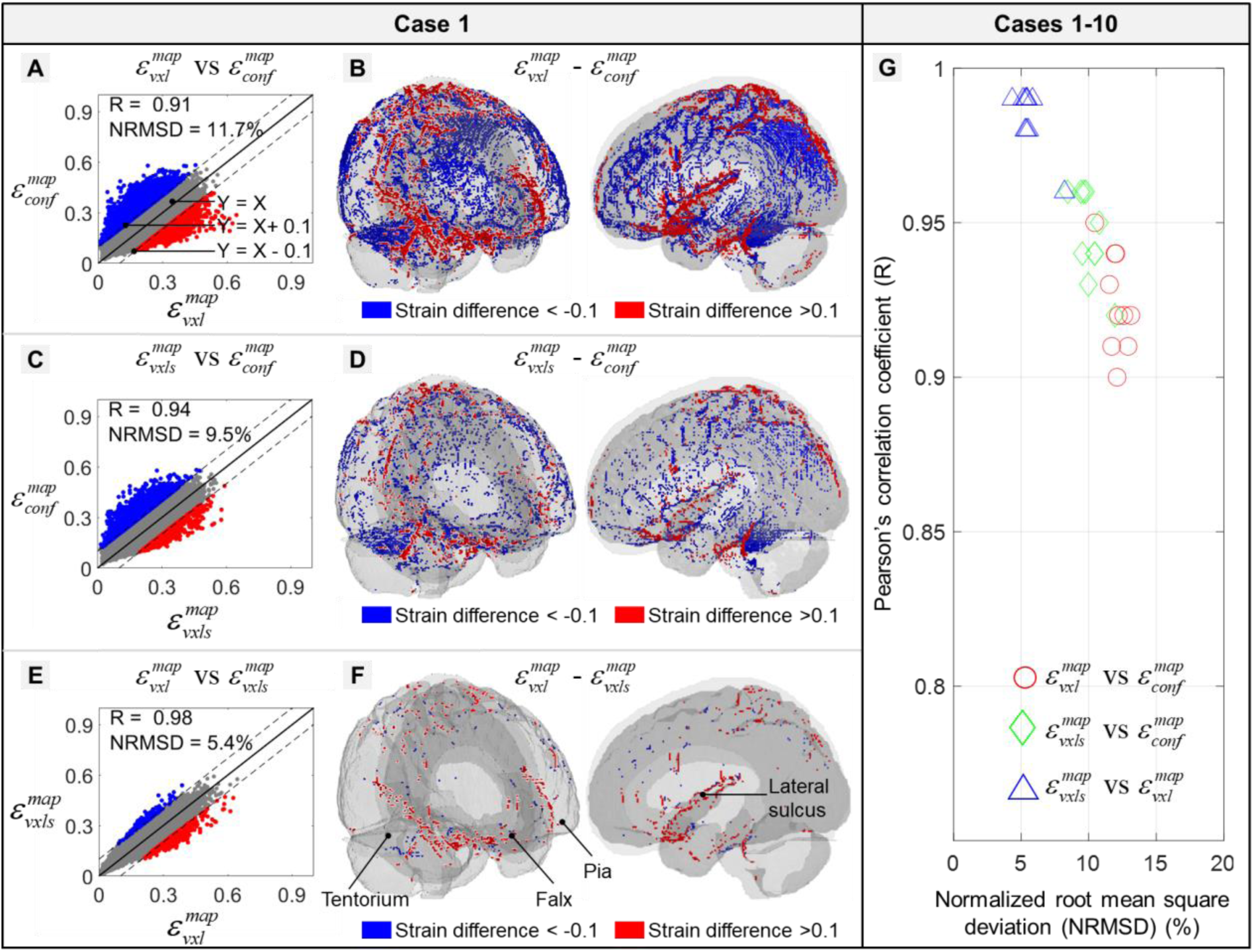
Comparisons of voxel-wise strain results among the Conf-, Vxl- and Vxls-models in Case 1 (A-F) and the Pearson’s correlation coefficient (R) and normalized root mean square deviation (NRMSD) in Cases 1-10 were summarized (G). (A) Scatter plots of voxel-wise strains estimated by Conf-model and Vxl-model. Those elements with high strain difference (i.e., the strain difference was either larger than 0.1 (in red) or less than -0.1 (in blue)) were plotted in isometric and side views in subfigure (B). To help the reader better perceive the location of these identified elements, the pia mater (light gray), falx, and tentorium (dark gray) were shown in half transparency in subfigure B. Similar plots were also presented between the Conf-model and Vxls-model in subfigures (C)-(D), and between the Vxls-model and Vxl-model in subfigures (E)-(F). Note that all the strain results were mapped to the ICBM space.

Results of voxel-wise strain comparisons in all ten cases were summarized in Fig 5G. Similar to the findings noted in Case 1, the largest strain difference was observed from the strain pair of *ε*_*conf*_*^map^* and *ε_vxl_^map^* (R: 0.90 ∼ 0.95, NRMSD: 10.5 ∼ 13.1%), followed by the pair of *ε*_*conf*_*^map^* and *ε _vxls_^map^* (R: 0.92 ∼ 0.96, NRMSD: 8.4 ∼ 11.2%). The smallest difference was noted between *ε _vxls_^map^* and *ε_vxl_ ^map^* (R: 0.96 ∼ 0.99, NRMSD: 4.3 ∼ 8.2%).

### Element quality

The Conf-, Vxl-, and Vxls-models varied in terms of element numbers and size (Table 2A). The Conf-model consisted of 4,612,951 solid elements (element size range: 0.1 ∼ 3.8 mm). The Vxl- and Vxls-models had the same element number (i.e., 2,116,684). For the element size, a uniform size of 1 mm was noted in the Vxl-model and a range of 0.4 mm to 1.3 mm was measured from the Vxls-model. The histogram of element size distributions was available in Appendix D.

**Table 2.** Element number, element quality, and volumes of key head components in the Conf-, Vxl-, and Vxls-models. (A) Element number and size range (in the unit of mm) of brain, subarachnoid cerebrospinal fluid (CSF), ventricle and whole head. (B) Element quality based on six indices, each of which the percentage of elements passing with criteria (e.g., 0.5 for Jacobian) and minimum value were listed. (C) Volume of brain, subarachnoid CSF, ventricle, and intracranial space.

| (A) | Element number (range of element size in the unit of mm) |  |  |  |  |  |  |  |  |  |  |  |
| --- | --- | --- | --- | --- | --- | --- | --- | --- | --- | --- | --- | --- |
| *Model* | *Brain* |  | *Subarachnoid CSF* |  | *Ventricle* |  | *Whole head* |  |  |  |  |  |
| Conf-model | 1,513,198 (0.1 ~ 3.8) |  | 1,483,377 (0.2 ~ 2.3) |  | 85,795 (0.2 ~ 1.9) |  | 4,612,951 (0.1 ~ 3.8) |  |  |  |  |  |
| Vxl-model | 1,567,589 (1) |  | 520,041 (1) |  | 29,054 (1) |  | 2,116,684 (1) |  |  |  |  |  |
| Vxls-model | 1,567,589 (0.4 ~ 1.3) |  | 520,041 (0.4 ~ 1.4) |  | 29,054 (0.4 ~ 1.4) |  | 2,116,684 (0.4 ~ 1.4) |  |  |  |  |  |
| (B) | Element quality index |  |  |  |  |  |  |  |  |  |  |  |
| *Model* | *Jacobian*<br>≥ 0.5 |  | *Warpage* (°)<br>≤ 30 |  | *Skew* (°)<br>≤ 60 |  | *Aspect ratio*<br>≤ 8 |  | *Min. angle* (°)<br>≥ 30 |  | *Max angle* (°)<br>≤ 150 |  |
|  | percent | Min. | percent | Max. | percent | Max. | percent | Max. | percent | Min. | percent | Max. |
| Conf-model | 98% | 0.45 | 92% | 111.8 | 99.9% | 70.0 | 99.9% | 6.6 | 99.8% | 18.0 | 99.9% | 161.9 |
| Vxl-model | 100% | 1 | 100% | 0 | 100% | 0 | 100% | 1 | 100% | 90 | 100% | 90 |
| Vxls-model | 91% | 0.13 | 87% | 83.5 | 100% | 42.5 | 100% | 3.0 | 100% | 44.7 | 97% | 158.1 |
| (C) | Volume of key head components (in the unit of mL) |  |  |  |  |  |  |  |  |  |  |  |
| *Model* | *Brain* |  | *Subarachnoid CSF* |  | *Ventricle* |  | *Intracranial space* |  |  |  |  |  |
| Conf-model | 1573 |  | 515 |  | 28 |  | 2116 |  |  |  |  |  |
| Vxl-model | 1568 |  | 520 |  | 29 |  | 2117 |  |  |  |  |  |
| Vxls-model | 1566 |  | 521 |  | 29 |  | 2116 |  |  |  |  |  |

The element quality of three models based on 6 indices was summarized in Table 2B. All elements in the Vxl-mode were perfect cubes and all quality indexes were fully scored (e.g., Jacobian: 1). The mesh smoothing procedure reduced the element quality in the Vxls-model. For example, the minimum Jacobian was 0.13 and only 91% of elements had a Jacobian over 0.5. When switching to surface-based meshing, Jacobian, skew, aspect ratio, and element angles fell within the desirable ranges, with over 98% of elements exceeding the criteria. For example, over 98% of elements in the Conf-model had a Jacobian over 0.5 with the minimum as 0.45. The only exception was noted in the warpage, in which only 92% of elements passed the criterion (i.e., warpage ≤ 30°) with the maximum as 111.8°. Detailed histograms of mesh quality were reported in Appendix D. The three models were highly similar in the size of the brain, subarachnoid CSF, ventricle, and intracranial space, as quantified by the cross-model volumetric differences of less than 5 mL (Table 2C).

## Discussion

The current study analysed the strain responses from three high-resolution, anatomically detailed head models, including one previously developed surface-based model with conforming elements for the brain-skull and brain-ventricle interfaces (Li et al. 2021) and two new models created from voxel-based meshing with and without smoothing. Other than the meshes, we kept all numerical settings in the three models exactly the same. When examining commonly used injury metrics, including the percentile strains below the maximum (e.g., 99 percentile strain) and volume fraction of brain elements with the strain over pre-defined thresholds, the responses of the three models were virtually equivalent. When scrutinizing the strain distribution, the three models showed different patterns at the interfacial boundary (e.g., sulci and gyri in the cortex, regions adjacent to the falx and tentorium) with strain differences over 0.1. Remarkably similar strain responses were noted at non-interfacial regions between the surface- and voxel-based models. The mesh smoothing procedure applied to the voxel-based model marginally reduced the strain discrepancies compared to the surface-based model. This study quantified the general similarity of strain responses between the surface- and voxel-based models and highlighted that caution should be exercised when interpreting the strain at the interface.

### Considerations of cross-model strain comparison

While comparing the strain responses among the three models, such as the non-100 percentile strain value (e.g., 99 percentile strain values to exclude the top 1 extremely deformed brain elements) (Fig 3) and volume fraction of brain elements with the strain magnitude over a certain threshold (Fig 4), virtually identical results were found between the surface- and voxel-based models with exceptions noted at the interfacial regions (Fig 5). Such findings correlated well with one early mouse study by Bradfield (2022), of which two FE head models of the mouse to virtually simulate experimental impacts. One mouse model possessed conforming meshes to capture the pial surface and the other was voxelized from the imaging segmentation without smoothing (analogous to the Conf-model and Vxl-model, respectively, developed in the current study). Similar to the current results, remarkably similar spatial strain distributions were noted between these two mouse models with some strain discontinuities noted at the brain cortex. Considering that the mouse is a lissencephalic species without the gyri and sulci, our study could be regarded as a significant extension to the previous effort by focusing on the human brain (gyrencephalic) with sophisticated integrations of the intricated foldings of the cerebral cortex and complex profiles of the ventricle systems. The general consistency between these two independent studies involving two species highlighted the credibility of our results.

The general similarity in brain strain response among the Conf-, Vxl-, and Vxls-models could be explained by their comparable element number for the brain as well as other identical numerical settings. For the brain mesh size, several previous studies highlighted its influence on the brain strain (Bradfield 2022; Giudice et al. 2019; Mao et al. 2013a; Panzer et al. 2013; Zhao and Ji 2019a). For example, Giudice et al. (2019) created a series of voxel models (without smoothing) and found reducing the element size from 4 mm to 1 mm increased the MPS (95^th^ percentile values) from 0.33 to 0.77. In light of this, the element number for the brain was deliberately ensured to be comparable among the three models (i.e., 1,513,198 for the Conf-model vs. 1,567,589 for the Vxl- and Vxls-models, Table 2A). Consequently, the average element sizes in the three models were all around 1 mm, although the exact distribution of element sizes was different due to inherent disparities in surface-based and voxel-based meshing techniques and the involvement of mesh smoothing (Appendix C). The three models were further harmonized in terms of other major determinants of brain responses, including geometric resources, material properties, element formulations, and interfacial conditions (Appendix A). Such a study design enabled a focused investigation on the influence of three rival meshing techniques on brain strain responses. Responses among the three models were extensively evaluated via qualitative visualization of strain contours (Fig 3A-I), quantitative metrics including cumulative percentile strain value (Fig 3J-L, Fig C1), cumulative volume fractions of the brain with MPS over different thresholds (Fig 4, Fig C2), and spatially detailed, voxel-wised strain comparisons (Fig 5). Our multi-aspect assessment protocol collectively verified the brain strain responses between surface- and voxel-based models were generally comparable.

The disparity in brain strain between the surface- and voxel-based models at the interfacial boundary was related to the geometric inaccuracies introduced by voxel-based meshing. Despite the three models being derived from the same imaging dataset, the surface-based model deployed conforming elements to precisely capture the input surfaces (i.e., brain, subarachnoid CSF, ventricles, and skull), while the voxel-based meshing, even with the smoothing operation, suffered from the intrinsic limitation of jagged interfaces between different tissues. Such a fundamental difference in resolving the geometrical complexity led to disparent strain responses at the interfacial region. Moreover, the mesh topologies between the surface- and voxel-based meshing were different (as exemplified by the enlarged views in Fig 1C vs. Fig 1D vs. Fig1E). Strain responses in the interfacial region with abrupt stiffness changes might be particularly sensitive to the localized mesh characteristics. For the brain-subarachnoid CSF interface, the brain was modelled as a hyper-viscoelastic material with a shear modulus of around 1 kPa, while the CSF as a nearly incompressible solid with low-shear resistance. For the falx-interhemispheric CSF-brain interfaces, the stiffness of the falx (Young’s modulus: ∼ 31.5 MPa) was much higher than those of the brain and CSF. When comparing the predicted strain between the Conf-model (conforming meshes for the cortical folds-subarachnoid CSF interface and brain-ventricle interface, Fig 1C) and Vxl-model (stair-stepped interface, Fig 1D), the brain elements with high strain difference were distributed at the site with shape stiffness gradient. The response dependency on the localized mesh topology at the region with material change has been previously reported in other TBI-related studies [e.g., cortical strain in the mouse brain (Bradfield 2022) and human brain (Giudice et al. 2019), pressure at the boundaries between different regions in blast-related head trauma (Yu and Ghajari 2019), etc], and even beyond the TBI context [e.g., strain energy density in a compressed metal plate with an air-filled hole (Guldberg et al. 1998), bladder deformation in response to the urinary pressure (Chai et al. 2012), etc].

The current study also found that the mesh smoothing marginally mitigated the strain difference between the surface- and voxel-based models. This was supported by the subtle variations in the R-values of voxel-wise strain correlations, i.e., 0.90 ∼ 0.95 between Conf-model and Vxl-model vs. 0.92 ∼ 0.96 between Conf-model and Vxls-model. Such a finding could be explained from two perspectives. First, although the mesh smoothing reduced the jaggedness level at the brain-CSF interface, the resultant profiles of the brain and ventricle in the Vxls-mode were still not identical to those in the Conf-model. Second, the mesh topology in the surface- and voxel-based meshing was disparate, which was speculated to be the dominant factor responsible for the strain differences between the Conf- and Vxls-models, as discussed above. This speculation was verified by the high correlations of the strain responses (R: 0.96 ∼ 0.99) between the Vxl- and Vxls-models, as both models shared identical mesh topologies.

The current study had important implications for the applicable scope of the voxel-based model. Due to the jagged interface, the voxel-based model was ubiquitously criticized in the literature [e.g., Giudice et al. (2019) and Li et al. (2021)]. However, such a criticism was often based on empirical speculation and has not been appropriately verified or falsified before. Our study filled this knowledge gap and found the strain responses of the surface- and voxel-based models were remarkably similar at the non-interfacial regions and substantially different at the interfacial regions. Such results thus supported the utilization of the voxel-based model to investigate brain responses at the non-interfacial regions and highlighted that caution should be exercised while interpreting the interfacial responses from the voxel-based model. For example, both Ho and Kleiven (2009) and Ghajari et al. (2017) used voxel-based head models to study brain trauma and found high strain at the depth of the sulci largely overlapped with the site of tau pathology in chronic traumatic encephalopathy. Despite interesting initial findings, our study highlighted that the conclusion from these two studies based on voxel-based models should be interpreted with care and further verifications were necessary by involving other advanced head models, e.g., the anatomically more authentic Conf-model (Li et al. 2021).

Desipte that the surface-based model was anatomically more authentic than the voxel-based models in terms of capturing the smoothing feature of the cortical folds-subarachnoid CSF and brain-ventricle interfaces of the human brain, no experimental data were available to support the investigation on which model had more biofidelic interfacial responses. As elaborated by Li et al. (2021) and Zhou et al. (2023), the responses of the Conf-model were validated by experimental data in terms of brain motion (Hardy et al. 2007), strain (Zhou et al. 2019b), and strain rate (Zhou et al. 2023) that were measured from sites away from the ventricles and cortex. This was also the case in other available experimental data (Alshareef et al. 2020; Dutrisac et al. 2021; Guettler et al. 2018; Hardy et al. 2001). In two promising exceptions, Mallory (2014) and Tesny (2022) used ultrasound transducers to measure the relative motion between the brain cortex and skull by imposing sagittal head rotations on cadaveric heads. However, the accuracy of reported motion data in these two studies seemed to be compromised by various compounding factors (e.g., post-mortem time, storage temperature, etc) (Mallory et al. 2024; Tesny 2022). Despite their uniqueness, these experimental datasets were not used to validate the cortical responses of FE head models in the current study. Another way of assessing the model-predicted results was to evaluate their predictability of injury. However, to the best of the authors’ knowledge, no real-life human injury dataset existed with definitive brain pathology at the interfacial boundaries paired with accurate measurement of impact kinematics. It was also worth mentioning that the aforementioned issue was not necessarily specific to the current study and, instead, represented a challenge in the TBI research field.

### Considerations of mesh techniques and element quality

For the Conf-model and Vxls-model in the curret study, the element quality was largely comparable to existing FE models (Garimella and Kraft 2017; Ho et al. 2009; Mao et al. 2013a; Trotta et al. 2020; Zhao and Ji 2019a; Zhou et al. 2016). For example, in one voxel-based head model (Ho et al. 2009) with mesh smoothing and the cortical folds and ventricles being preserved, the most distorted element has a Jabocian of 0.23, inferior to the Conf-model (minimum Jacobian: 0.45) and superior to the Vxls-model (minimum Jacobian: 0.13). In another FE head model (average element size: ∼ 1.1 mm) without the gyri and sulci (Zhao and Ji 2019a), the minimum and maximum angles were 11.8° and 176.3°, in comparison to 18.0° and 161.9° in the Conf-model, 44.7° and 158.1° in the Vxls-model. For the Vxl-model developed in the current study and its other counterparts in the literature (Alshareef et al. 2021; Giudice et al. 2020; Miller et al. 2016; Pavan et al. 2022), all elements were perfect cubes with idealized mesh qualities.

For the surface-based meshing, to the best of the authors’ knowledge, only the Conf-model (Li et al. 2021) and a few other models developed by the same research group (Li et al. 2021; Zhou et al. 2022a; Zhou et al. 2020a) employed pure hexahedral meshes to discretize the whole head with the cortical folds and ventricles preserved. For the voxel-based meshing without smoothing, it has been leveraged by several groups to develop head models with stair-like interfaces (Alshareef et al. 2021; Giudice et al. 2020; Miller et al. 2016; Pavan et al. 2022). When it came to the mesh smoothing, the choice of one Laplacian-based, volume-preserving meshing algorithm in the current study was justified by the availability of the technique details elaborated by Boyd and Müller (2006) and its early implementation to smooth voxel-based FE models for brain research (Chen and Ostoja-Starzewski 2010; Ho and Kleiven 2009). In the study by Chen and Ostoja-Starzewski (2010) in which the mesh in a voxelized whole-head model was smoothed, element quality statistics were not reported, prohibiting a direct comparison to the Vxls-model in the current study. In the other study by Kelley et al. (2015), the mesh smoothing was only applied to brain sections (i.e., partitioning the brain into eight sections with the size of each section around one-eighth of the whole brain), instead of the whole-head level as was the case of the Vxls-model in the current study. It was also noted that the exact details of the smoothing procedure were omitted in a few studies (Duckworth et al. 2022; Griffiths et al. 2023) while developing voxel-based FE human head models.

### Other limitations and paths forward

Several other limitations, except for the few highlighted above, should be acknowledged with further investigations deemed necessary. First, the current study approximated the brain as a homogeneous structure, as no consensus has been reached yet regarding the regional trend of brain stiffness which seemed to be compounded by various factors (e.g., loading rate, drainage conditions, length scale, etc) (Budday et al. 2020). Future work could plant region-wise material properties, when more consensual results were avaialble, to the FE head model with meshes conforming to the material boundaries (Zhang et al. 2010). Second, the CSF was simplified as a Lagrangian-based solid structure with low shear stiffness, as was also the case in other FE head models (Alshareef et al. 2021; Chen and Ostoja-Starzewski 2010; Duckworth et al. 2022; Giudice et al. 2020; Giudice et al. 2019; Griffiths et al. 2023; Ho and Kleiven 2009; Ho et al. 2009; Ho et al. 2017; Madhukar and Ostoja-Starzewski 2019; Miller et al. 2016; Pavan et al. 2022). Emerging efforts have been noted in implementing arbitrary Lagrangian-Eulerian (ALE) fluid elements for the ventricle (Zhou et al. 2022a; Zhou et al. 2020a). Such an implementation could be expanded to subarachnoid CSF, although the enormously high computational expense per solving the ALE formulation (Souli et al. 2000) was expected to be a bottleneck. Beyond the three rival techniques presented in this study, extra hexahedral meshing approaches were available, e.g., the multi-block approach that was prevalently used in the biomechanical field (Alvarez and Kleiven 2018; Bradfield 2022; Garimella and Kraft 2017; Giordano and Kleiven 2016; Grosland et al. 2009; Kallemeyn et al. 2009; Mao et al. 2013a; Schonning et al. 2009; Zhao and Ji 2019a; Zhou et al. 2016). However, the multi-block approach has not succeeded yet in developing hexahedral-based head models with details such as gyri and sulci retained and hence were not included in the current study. The current study employed the Lagrangian FE method, wherein the element follows the material deformation, making the development of a high-quality hexahedral mesh essential for numerical stability and reliability. As extensively reviewed by Wittek et al. (2016), alternative meshless methods, e.g., material point method (Lu et al. 2019), smoothed-particle hydrodynamics method (Duckworth et al. 2021), incompressible computational fluid dynamics method (Atsumi et al. 2021; Miller et al. 2012; Zhang et al. 2013), could circumvent excessive mesh distortion in the Lagrangian FE method. For example, Miller et al. (2012) and Zhang et al. (2013) implemented a Meshless Total Lagrangian Explicit Dynamics method to simulate the craniotomy-induced brain shift and thus did not confront the challenge of developing high-quality mesh, as investigated in the current study. Lastly, the three models used in the current study contain millions of elements. Due to the extensive computational time and substantial resource demands, these models were practically unsuitable to perform large-scale impact simulations. As inspired by the works by Zhan et al. (2024) and Wu et al. (2022), machine learning techniques could be employed to capture the complex, nonlinear relationship between impact kinematics and brain strain predicted by high-resolution FE head models. Such a machine learning model could rapidly and reliably predict brain strain under known loading conditions, eliminating the need for time-consuming and resource-intensive simulations.

## Conclusion

The current study performed strain comparisons among three anatomically detailed FE head model models. One was previously developed from surface-based meshing with conforming elements for the cortical folds-subarachnoid CSF and brain-ventricle interfaces (Li et al. 2021), while the other two were newly created by converting the segmented voxels into elements with and without mesh smoothing. Except for the meshes, other numerical settings were the same among the three models. It was verified that the strain responses between the surface- and voxel-based models were largely comparable with exceptions at the interfacial region (e.g., sulci and gyri in the cortex, regions adjacent to the falx and tentorium) where strain differences over 0.1 were noted. By scrutinizing the commonly used injury metrics, including the percentile strain below the maximum (e.g., 99 percentile strain) and the volume fraction of brain elements with strains over certain thresholds, the responses of the three models were virtually identical. The smoothing procedure marginally mitigated the strain differences between the surface- and voxel-based models. This study provided quantitative insights into how close the strain responses were between the surface- and voxel-based FE head models and highlighted the importance of exercising caution when using strain at the interface to predict injury.

## Acknowledgement

This research has received funding from KTH Royal Institute of Technology (Stockholm, Sweden), the Swedish Governmental Agency for Innovation Systems (VINNOVA) (no. 2023-00753 and no. 2024-03635), Swedish Research Council (VR-2020-04496, VR-2020-04724, and VR-2024-02782), Skyltfonden from the trafiksäkerhet (TRV 2024/23811 and TRV 2024/50301), Carl Tryggers Foundation (CTS 24: 3284), and the Torvald and Britta Gahlin’s foundation. The content of this article is solely the responsibility of the authors and does not necessarily represent the official views of neither funding agencies. The computational simulations were enabled by resources provided by the National Academic Infrastructure for Supercomputing in Sweden (NAISS) at the center for High Performance Computing (PDC) partially funded by the Swedish Research Council through grant agreement no. 2022-06725. The authors declare that they have no known competing financial interests or personal relationships that could have appeared to influence the work reported in this paper.

## Appendix A. Geometry, material, interface, and element formulation of surface- and voxel-based models

**Table A1.** Commonalities in terms of geometric sources, material properties, element formulation, and interfacial conditions in the Conf-model, Vxl-model and Vxls-model

| **Geometry** | **Resource** |
| --- | --- |
| Brain/ventricle/subarachnoid<br>CSF/skull | ICBM 152 atlas |
| **Material** | **Description** |
| Brain | Ogden hyperelastic and linear viscoelastic constants: $\mu_1=53.8$ Pa, $\alpha_1=10.1$, $\mu_2=-120.4$ Pa, $\alpha_2=-12.9$; $G_1=0.31$ MPa, $G_2=78$ kPa, $G_3=6.2$ kPa, $G_4=8.0$ kPa, $G_5=1.0$ kPa, $G_6=3.0$ kPa; $\beta_1 \dots \beta_6 = 10^6 \dots 10^1$ (1/s), where $\mu_i$ and $\alpha_i$ are Ogden parameters, $G_i$ represents the shear relaxation module, and $\beta_i$ represents the decay constants |
| Cerebrospinal fluid | Elastic fluid with the bulk modulus as 2.1 GPa |
| Falx/tentorium/dura mater | Simplified rubber with experimentally derived stress-strain curve |
| Pia mater | Simplified rubber with experimentally derived stress-strain curve |
| Skull | Rigid |
| **Element** | **Description** |
| Brain/cerebrospinal fluid/skull | Hexahedral elements with selectively reduced integration (ELFORM = 2) |
| Falx/tentorium/dura mater/pia mater | Quadrilaterals with fully integrated Belytschoko-Tsay membrane (ELFORM = 9) |
| **Interface** | **Description** |
| Brain-subarachnoid cerebrospinal fluid interface | All interfaces shared common nodes, and no contact algorithm was defined. |
| Subarachnoid cerebrospinal fluid-skull interface |  |
| Brain-ventricle |  |

## Appendix B. Strain of the surface-based model and its counterparts mapped in the image space

**Fig B1.**
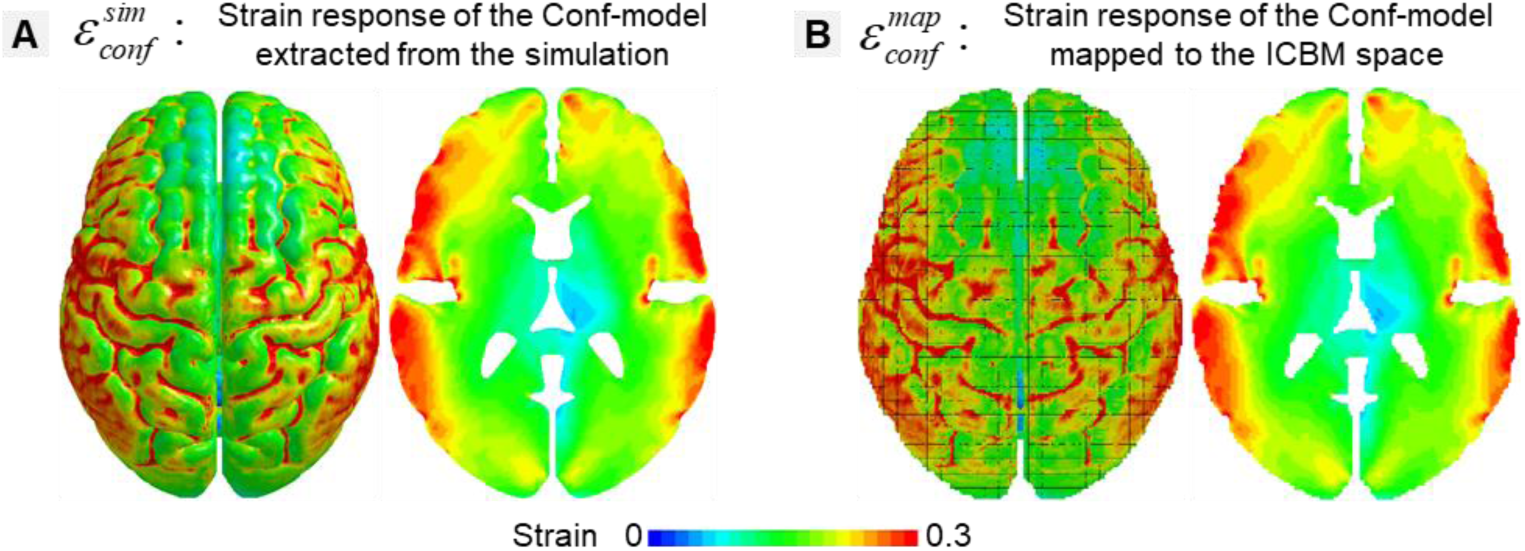
(**A**) Element-wise strain peaks directly simulated by the Conf-model (i.e., *ε_conf_^sim^*). (**B**) Voxel-wise strain peak by mapping strain peaks to the ICBM spaces (i.e., *ε_conf_^map^* ). Visually identical strain distribution was noted before and after the mapping, indicating the voxelization procedures captured the strain-related feature of the Conf-model.

## Appendix C. Cumulative curves of percentile strains and volume fractions estimated for Surf- and Voxel-based models for Cases 2-10

**Fig C1.**
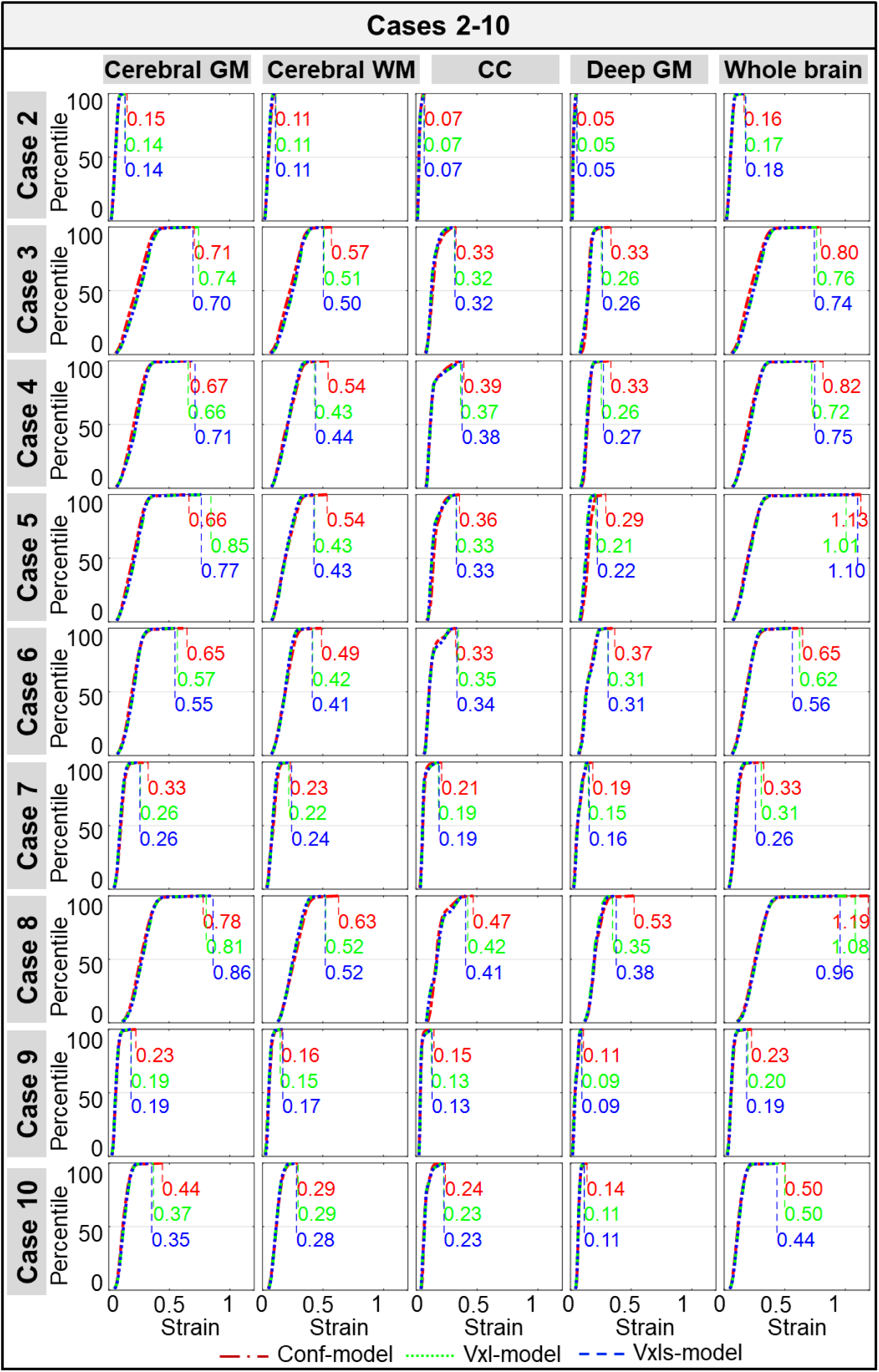
Cumulative percentile strain curves estimated by the Conf-model, Vxl-model, and Vxls-model in Cases 2-10 with the 100th percentile strain peaks listed. These curves were calculated in the cerebral gray matter (GM), cerebral white matter (WM), corpus callosum (CC), deep GM, and whole brain.

**Fig C2.**
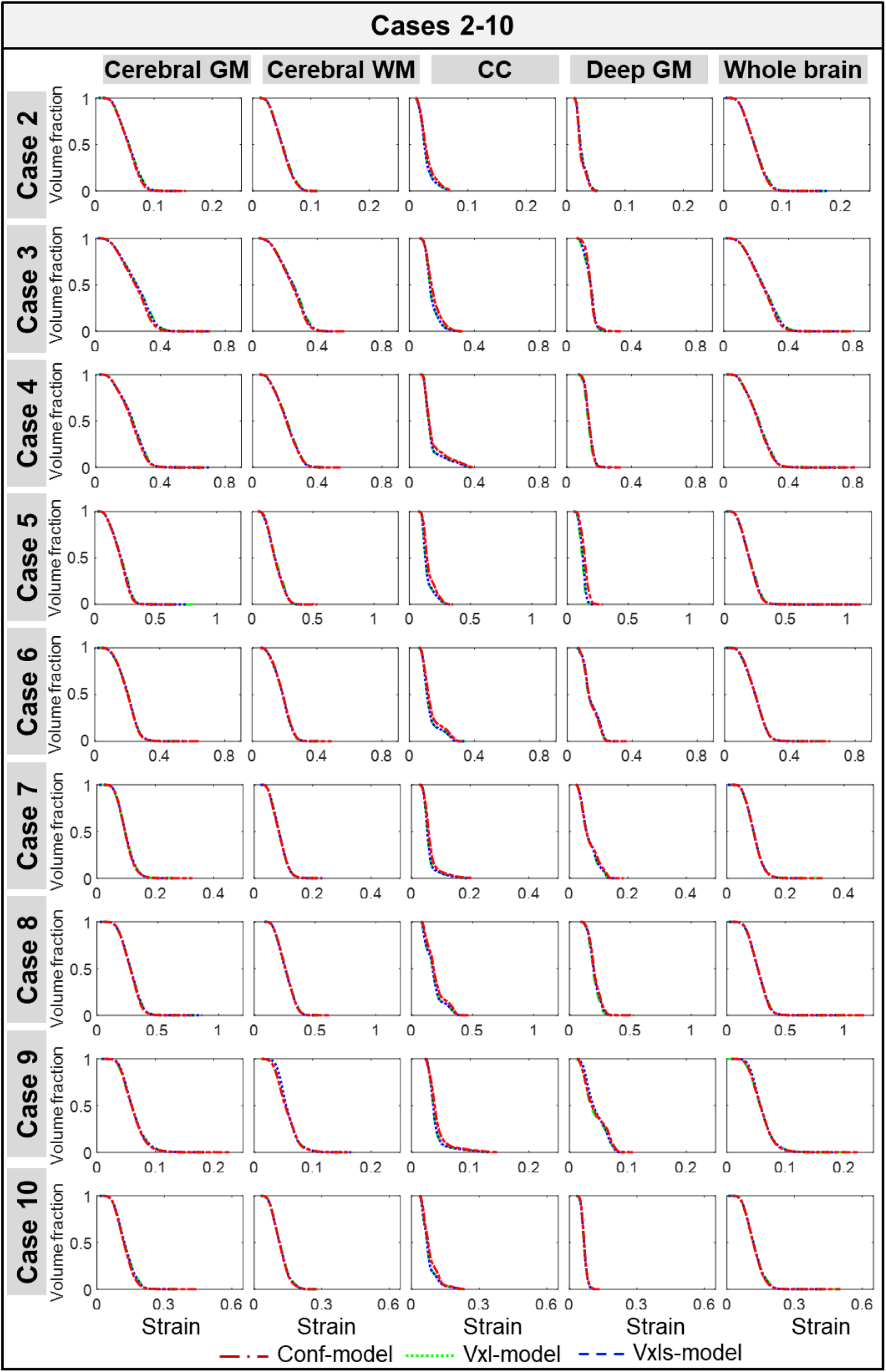
Cumulative curves of volume fraction of the elements with the strain over varying thresholds estimated by the Conf-model, Vxl-model, and Vxls-model in Cases 2-10. These curves were calculated in the cerebral gray matter (GM), cerebral white matter (WM), corpus callosum (CC), deep GM, and whole brain.

## Appendix D. Histograms of element number, element size, and mesh quality

**Fig D1.**
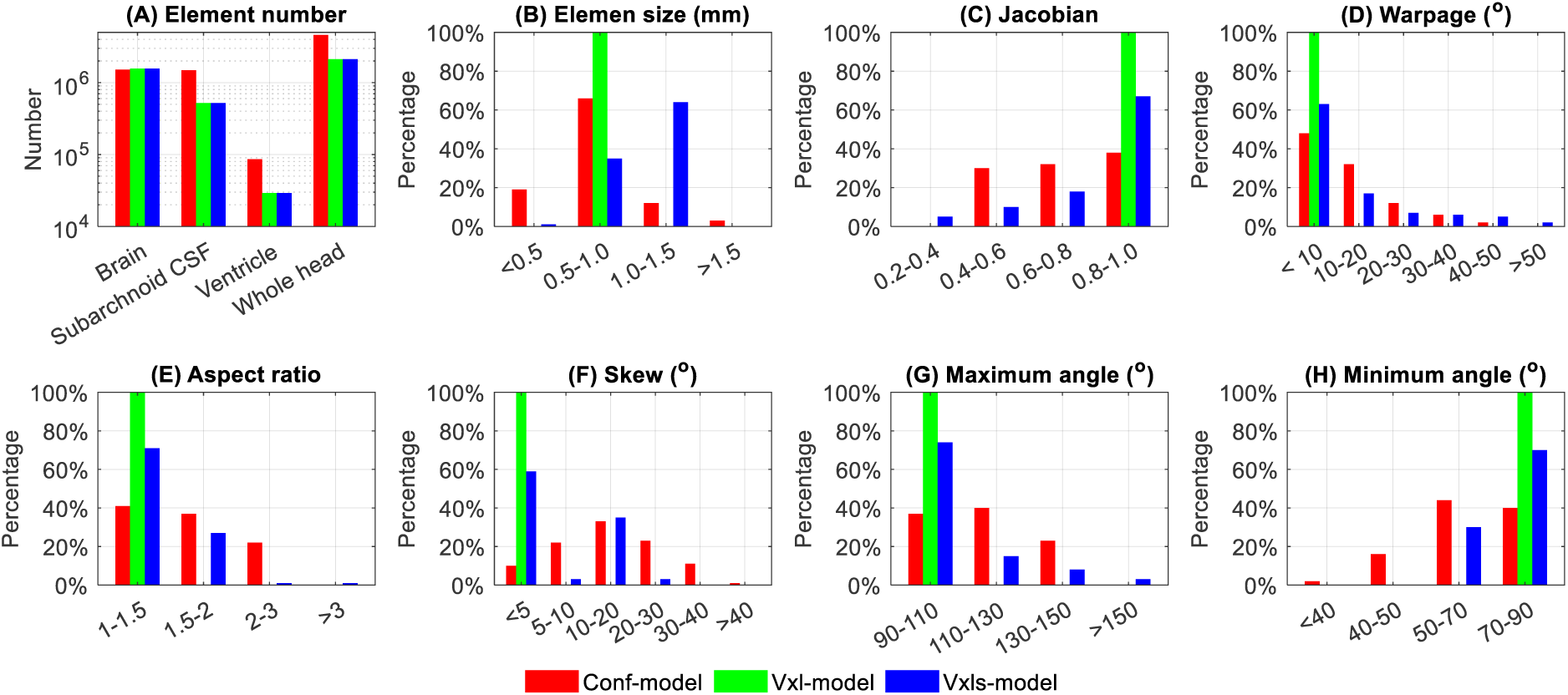
Histograms of element number (A), element size (B), and mesh quality based on 6 indices (C-H) of the Conf-model, Vxl-model, and Vxls-model.

